# Heterogenous Biofilm Mass-Transport Model Replicates Periphery Sequestration of Antibiotics in *P. aeruginosa* PAO1 Microcolonies

**DOI:** 10.1101/2023.07.28.551018

**Authors:** Joshua Prince, A-Andrew D. Jones

## Abstract

A spatiotemporal model for antibiotic accumulation in bacterial biofilm microcolonies which leverages heterogenous porosity and attachment site profiles replicated the periphery sequestration phenomena reported in prior experimental studies on *Pseudomonas aeruginosa PAO1* biofilm cell clusters. These *P. aeruginosa* cell clusters are *in vitro* models of the chronic *P. aeruginosa* infections found in adult cystic fibrosis patients, which display resistance to antibiotic treatments, leading to exacerbated morbidity and mortality. This resistance has been partially attributed to periphery sequestration, where antibiotics are unable to penetrate biofilm cell clusters. The underlying physical phenomena driving this periphery sequestration have not been definitively established. This paper introduces mathematical models to account for two proposed physical phenomena driving periphery sequestration: biofilm matrix attachment and volume-exclusion due to variable biofilm porosity. An antibiotic accumulation model which incorporated these phenomena was able to better fit observed periphery sequestration data compared to previous models.

## 2 Introduction

Cystic fibrosis is a genetic disorder mainly caused by mutation in gene for the CFTR protein with a patient population of around 100,000 worldwide (1). People with cystic fibrosis in the United States have a median lifespan of 48.4 years (2). Recently, the drug class known as CFTR-modulators have significantly improved outcomes for people with cystic fibrosis (1). However, even patients using CFTR-modulators acquire chronic *Pseudomonas aeruginosa* infections (1). In most cases of chronic *Pseudomonas aeruginosa* lung infections, the bacteria have formed a biofilm that is recalcitrant to common antibiotic therapies (3). Two classes of hypothesis have formed to explain this antibiotic-biofilm recalcitrance: antibiotic diffusion-limitations and physiology-based mechanisms (4). The diffusion-limitations hypothesis is that specific or non-specific physical interactions between an antibiotic and the biofilm extra-cellular matrix lead to slower antibiotic penetration into biofilms, preventing antibiotics from reaching and killing interior cells (5). Mathematical models based on this hypothesis quantify this diffusion-limitation by assuming a biofilm with homogenous porosity and antibiotic attachment site density and incorporating antibiotic-matrix interactions into a “homogenized” diffusivity term (6, 7). Experimental studies however have indicated that antibiotic-matrix interactions primarily cause biofilm recalcitrance not due to hindered diffusion but due to spatial heterogeneity in the biofilm causing antibiotics to accumulate at the periphery of the biofilm at equilibrium, termed “periphery sequestration” (8, 9). To account for this periphery sequestration theory of biofilm recalcitrance, we developed a mathematical model that could relax either or both the assumptions of homogenous biofilm porosity and antibiotic attachment site concentration used in prior mathematical models (10, 11). Only the mathematical model which accounted for heterogeneities in both antibiotic attachment sites and porosity replicated the antibiotic accumulation of both ciprofloxacin and tobramycin in *P. aeruginosa* cell-clusters determined experimentally by Tseng *et al* (9). This new physical model for antibiotic accumulation in biofilms undergirds a new conceptual model of how antibiotic-matrix interactions lead to biofilm recalcitrance. These new models can be applied in designing new antibiotics for chronic *P. aeruginosa* infections for people with cystic fibrosis to circumvent the periphery sequestration mechanism of biofilm-recalcitrance. Further the methods developed here to capture heterogeneities in both attachment sites and porosity could be applied to transport limitations through other biological hydrogels, like mucosal membranes (5).

## 3 Theory

Accumulation of antibiotics into bacterial biofilm microcolonies was modelled using a diffusion-reaction mass-transport approach (12, 13). The biofilm was approximated as a thin-film of height *H* [m] along a single coordinate axis of depth represented by *x*. The system boundaries were a fluid-biofilm interface and a solid-biofilm interface, both normal to the depth axis of the biofilm and parallel to each other. The liquid-biofilm interface was a constant source of antibiotics *c*_0,*I*_,[kg/m^3^] . The solid-biofilm interface was an antibiotic-impermeable substratum, defined as the origin of the one-coordinate system (x = 0). Antibiotic molecules in the biofilm were defined as either in the biofilm pore space and mobile (*m*_M_[kg]) or attached to biofilm biomass (*m*_A_[kg]). Antibiotic concentrations were defined on eithera total biofilm volume basis, *C*_*M,T*_ = *m*_*M*_/*V*_*T*_[kg m^3^], or a biofilm interstitial-volume basis, *C*_*M,I*_ = *m*_*M*_/*V*_*I*_[kg m^3^]. The antibiotic was assumed to be cell impermeable. Biofilm porosity was defined as the ratio of interstitial biofilm volume to total biofilm volume, *φ* = *V*_*I*_/*V*_*T*_, and was used to relate the two concentration definitions, as *C*_M,T_ =*φC*_*M,I*_ and *C*_*A,T*_*=φC*_*A,I*_.Using these definitions, the dimensionless mass-balances on mobile, *Ĉ*_M_ = *C*_*M,I*_/*C*_0,*I*_,and attached antibiotic, *Ĉ*_*A*_ = *C*_*A,I*_/*C*_0,*I*_, respectively, were derived using the Fick-Jacobs expression (14) for diffusion in systems with varying cross-section and an assumption of reversible antibiotic attachment as follows (see Supplementary Info):

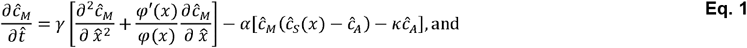

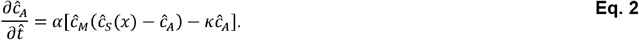

Dimensionless time, 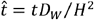, and position, 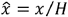, were used to non-dimensionalize, where *D*_*W*_[*m*^2^/*s*] is the antibiotic diffusivity in water. The dimensionless groups which arose from this non-dimensionalization included: *γ* = *D*_*B*_*/D*_*w*_, the effective diffusivity of the antibiotic in the biofilm; *κ* = *K*_*M,I*_*/K*_*A,I*_*C*_*O,I*_, the dimensionless attachment equilibrium constant where *K*_*A*/*M,I*_ are the rate of antibiotic attachment/detachment; and *α* = *K*_*A,I*_*C*_*O,I*_*H*^2^/*D*_*w*_, the dimensionless ratio of antibiotic attachment rate to diffusion, similar to a Thiele Modulus. The variable cross-sectional area available for diffusion from the Fick-Jacobs expression is accounted for by the porosity factors in the diffusion term (see derivation in SI). To differentiate this model from previous homogenous biofilm models, attachment site concentration and porosity were assumed to be heterogenous throughout the depth of the biofilm. The dimensionless attachment site concentration, *Ĉ* = *C*_*S,I*_/*C*_*O,I*_, profile was assumed to take the form 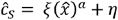, where *ξ =* (*C*_*S,LI*_ – *C*_*S,SI*_)/C_*O,I*_,was the dimensionless attachment site heterogeneity constant, *η* = *C*_*S,SI*_/*C*_*O,I*_, was the dimensionless solid-interface attachment site concentration, *C*_*S,LI*_, [# sites/m^3^] was the liquid-biofilm interface attachment site concentration, *C*_*S,SI*_,[# sites/m^3^] was the solid-biofilm interface attachment site concentration, and was a profile shape parameter. The porosity profile was assumed to take the form 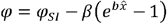, where *φ*_*SI*_ was the porosity at the solid-biofilm interface, *β* = (*φ*_*SI*_ - *φ*_*LI*_)/(*e*^*b*^ −1) was the porosity heterogeneity constant, *φ*_*LI*_ was the porosity at the liquid-biofilm interface, and *b* was a profile shape parameter. These attachment site and porosity profiles are not assumed to be universal across antibiotic-bacterial systems, as evidenced in other literature (15). Spatial concentration profiles for *Ĉ*_*M*_ and *Ĉ*_*A*_ calculated by solving Eq. 1 and Eq. 2 using a set of dimensionless parameter values were translated to total concentration profiles on a total biofilm-volume basis using the definition *ĈT,T = φ* (*Ĉ*_*M,I*_ *+ Ĉ*_*A,I*_) To quantify the degree of antibiotic sequestration with the biofilm, a “degree of sequestration” quantity, *Se*, was defined quantitatively as

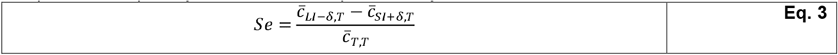

where 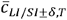 was the equilibrium average total dimensionless concentration on a total volume basis within 5% of the liquid/solid-interface, respectively. A positive value of was associated with antibiotic periphery sequestration, while a negative value *Se*of was associated with the antibiotic interior sequestration.

## 4 Results and Discussion

This proposed heterogenous biofilm model was tested for its ability to predict periphery sequestration of antibiotic within the biofilm at equilibrium. The effects of the two biofilm heterogeneity constants, *ξ* ∈ [-10,10] and *β*^*\**^ *= φ*_*S,I*_ - *φ*_*L,I*_∈[−0.5,0.5], on *Se* show that positive *ξ* and negative *β* were associated with periphery sequestration, and vice-versa for interior sequestration (Fig. 1). A positive *ξ* was attributed to a relatively large amount of attachment sites for the antibiotic near the periphery to sequester at compared to the interior. This highlights the divergence of this model from the common Crank assumption set for antibiotic-biofilm accumulation and allows for a depth-dependent attachment site concentration site profile (5). A negative *β* leading to periphery sequestration was attributed to a relatively large amount of pore space available for the antibiotic to occupy near the biofilm periphery compared to the interior. This highlights the models ability to account for a “volume-exclusion” effect of porosity where antibiotic is unable to occupy cell-volume, distinguishing it from the Hinson and Kocher method of porosity incorporation, which only modifies the diffusion constant (11). Both the homogenous biofilm model (10) and the homogenized biofilm model (6) can be represented as the origin of the investigated phase-space and shows no degree of sequestration, highlighting their inability to predict periphery sequestration at equilibrium.

**Figure 1.**
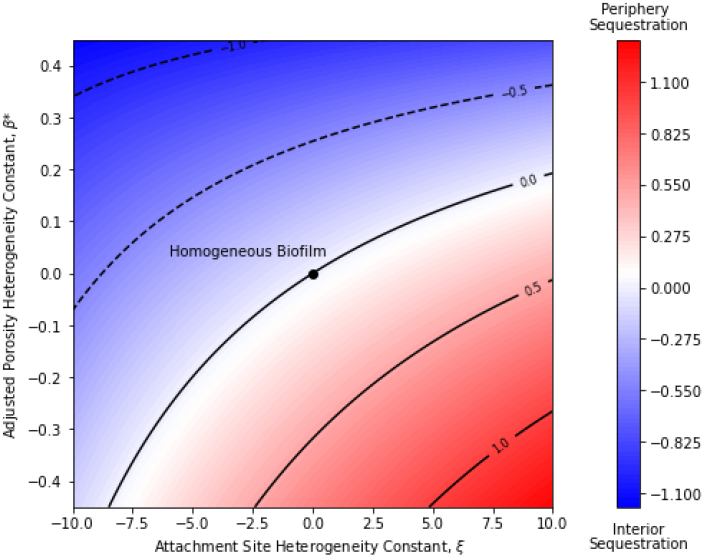
Phase-space plot showing effect of adjusted porosity and attachment site heterogeneity constants (*β*^*\**^and *ξ*, respectively) on degree of antibiotic sequestration within the biofilm, *Se*. Positive *Se* correspond to periphery sequestration (red), negative *Se* correspond to interior sequestration (blue). Contour lines of *Se* shown along with heatmap. Parameter values of *γ* = 1, *α* =1, *b* = 5, *a* = 5, *η* = 15, *φ*_*SI*_ = 0.5, and *k* = 1 used for solving Eq. 1 & 2 with given, *β*^*\**^ and *ξ* values.

The proposed heterogenous biofilm model replicates the periphery sequestration of tobramycin and ciprofloxacin in *P. aeruginosa* PAO1 biofilm microcolonies dynamically, with experimental data acquired from Tseng *et al* (Fig. 2). Two goodness-of-fit criteria to quantify model fit, the residual sum of squares, RSS = ∑_*i*_*y*_*o,i*_ – *y*_*p,i*_and the Akaike Information Criteria, *AIC* = 2*p* + *N* [*ln*(*2π* ∑_*i*_ *RSS*/*N*) + 1], where is the number of model parameters and is the number of data-points, were calculated for each model fit to the data. Fits were determined by systematically varying parameter values to find local RSS minimums for each parameter manually. The use of the model with the assumption of a homogenous biofilm led to a poor fit to the observed accumulation data for both antibiotics (Fig. 2A,B). A similarly poor fit was found for a model with a heterogenous porosity profile with homogenous attachment site profile (Fig. 2C,D). The heterogenous attachment site profile with a homogenous porosity profile replicated the tobramycin accumulation data (Fig. 2E) but had poor goodness-of-fit measures to the ciprofloxacin data (Figure 2F). The model with both a heterogeneous porosity profile and heterogenous attachment site profile (Fig. 2G,H) was able to replicate both the tobramycin and ciprofloxacin accumulation, with the AIC goodness-of-fit criteria, which accounts for number of model parameters, as low or lower than all other models for both antibiotics (Fig. 2I). This is primarily because of the full model’s ability to replicate the maximal concentration seen near the biofilm periphery. Concentrations at early time-points are likely underestimated due to implementation of the proposed model in Cartesian coordinates rather than the cylindrical coordinates more appropriate for this experimental system. Future theoretical work could implement algorithmic RSS-minimization techniques for model fits to ensure global minimums over a domain of interest are found. Future experimental work could spatiotemporally measure antibiotic accumulation in biofilm microcolonies concurrently with biofilm porosity profiles and test if the latter improves predictions of the former using the proposed model.

**Figure 2.**
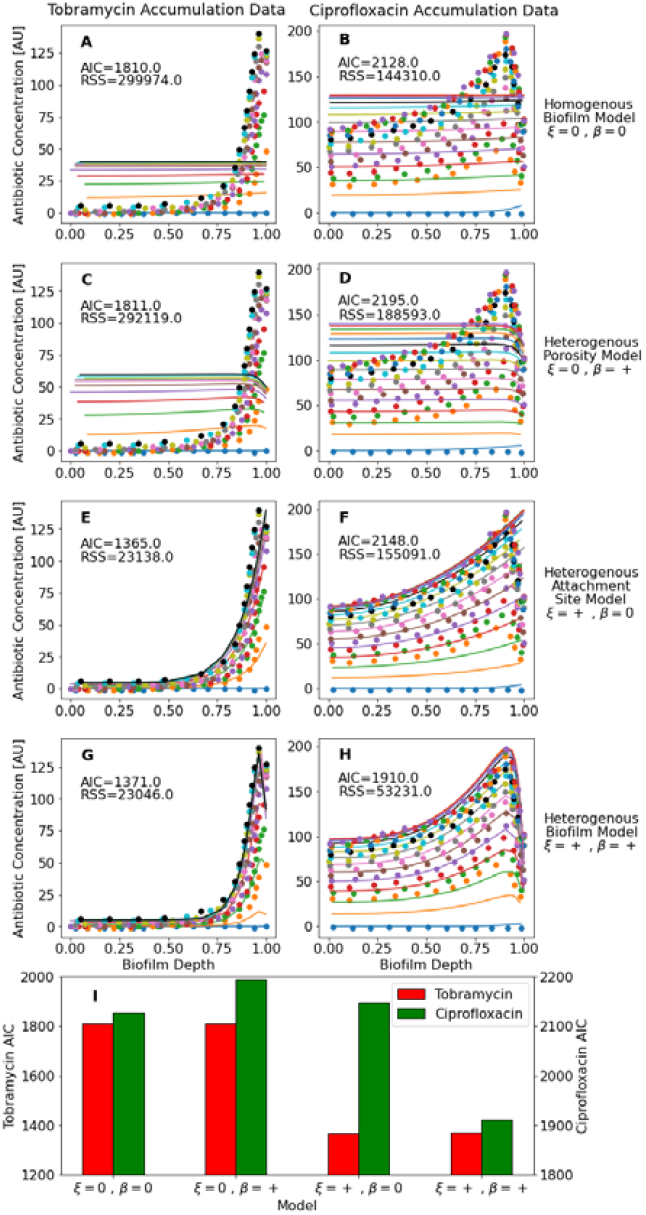
Fits of homogenous biofilm model (A,B), heterogenous porosity model (C,D), heterogenous attachment site model (E,F), and full heterogenous biofilm model (G,H) to data on accumulation of tobramycin (A,C,E,G) and ciprofloxacin (B,D,F,H) into *P. aeruginosa* PAO1 microcolonies. Dots represent accumulation data from Figure 2 of Tseng *et al*, with different colors representing different time-points. Solid lines represent fits of the data using the respective model, with color matched between literature data and model fits. An assumption of negative *β* for the porosity profile was inferred from biofilm cross-sections where antibiotic accumulation was tracked. (I) Comparison of AIC goodness-of-fit criteria of each model to the tobramycin and ciprofloxacin datasets. Parameter values for fits included in SI.

## 5 Methods

To solve the steady-state form of governing equations used in this paper for equilibrium concentration profiles, a finite-differences solver based on spatial discretization along with the Newton-Raphson technique for the resulting set of non-linear equations was implemented in python in the SPYDER IDE, with code available on Github (https://rb.gy/o8x7v). To solve the time-dependent governing equations for dynamic concentration profiles, the method-of-lines technique was incorporated into the same solver system. Additionally, a variable liquid-interface concentration boundary replaced the constant concentration BC for this dynamic solver due to the observed dynamic BC in the literature data. Relative antibiotic concentrations were inferred from Tseng *et al* using WebPlotDigitizer (9).

## Supporting information

Supplmental Equation Derivation

## Data, Materials, and Software Availability

All study data are included in the article, SI Appendix or GitHub repository (https://rb.gy/o8x7v).

## 6 Acknowledgements

Research reported in this publication was supported by the National Institute of General Medical Sciences of the National Institutes of Health under Award Number NIH R35 GM142898. The content is solely the responsibility of the authors and does not necessarily represent the official views of the National Institutes of Health. The authors would like to thank Phil Stewart at Montana State University for helpful discussions on prior models.

## Author contributions

J.P. conceptualized the model, and implemented the model numerically, and performed the data analysis. The original draft was written by J.P. Review and editing of the draft were carried out by J.P. and A.J. Project conception, administration and supervision was the responsibility of A.J.

## Competing interests

All authors declare no competing interests.

## Notes

https://rb.gy/o8x7v

### Competing Interest Statement

The authors have declared no competing interest.

