## Supplementary material for "Heterogenous Biofilm Mass-Transport Model Replicates Periphery Sequestration of Antibiotics in *P. aeruginosa* PAO1 Microcolonies": Supplmental Equation Derivation

Joshua Prince^1,2^ and A-Andrew D. Jones, III^1,2,3^

1. Department of Civil and Environmental Engineering, Duke University
2. Integrated Toxicology and Environmental Health Program, Duke University
3. Thomas Lord Department of Mechanical Engineering & Materials Science

A-Andrew D. Jones, III

**This PDF file includes:**

Supporting text

SI References

**Derivation of Governing Equation.**

The derivation of the mass-balances for dimensionless mobile and attached antibiotic starts with taking a control-volume within the biofilm. The control-volume is a thin cross-section of the biofilm starting at depth $x_{1}$ in the biofilm with thickness $\Delta x$ and cross-sectional area $A_{T}$, giving it a volume $V_{T}=$ $\Delta xA_{T}$. The control volume is composed of bacterial cells and interstitial space. The antibiotic is assumed to be cell-impermeable, which leads to the antibiotic only occupying the interstitial space. The total mass of antibiotic in both the total control-volume and interstitial space $m$. The antibiotic which is mobile and within the pore space is defined as $m_{M}$, while the antibiotic in the interstitial space and attached to biofilm biomass is defined as $m_{A}$. A mass-balance on both of these conserved quantities can be taken within the control-volume, yielding

| $\frac{dm_{M}}{dt}=\dot{m}_{M,x}-\dot{m}_{M,x+\Delta x}-\dot{m}_{M,att}+\dot{m}_{M,mob}$ and | Eq. S1 |
| --- | --- |
| $\frac{dm_{A}}{dt}=\dot{m}_{M,att}-\dot{m}_{M,mob}$, | Eq. S2 |

where $\dot{m}_{M,x_{1}/x_{1}+\Delta x}$ represent the rate of mobile antibiotic mass entering/exiting the control-volume, $\dot{m}_{M,att}$ represents the rate of antibiotic attaching the biofilm biomass, and $\dot{m}_{M,mob}$ represents the rate of antibiotic mobilizing from the biofilm biomass. The rates of mobile antibiotic exiting and entering the control-volume can be written in terms of flux using the expression $\dot{m}_{M,x_{1}/x_{1}+\Delta x}=\left( J_{T}A_{T} \right)_{x_{1}/x_{1}+\Delta x}$, where $A_{T}$ is the total cross-sectional area of the biofilm and $J_{T}$ is the flux through that area. However, since the antibiotic is only mobile in the interstitial volume, this can be equivalently written as $\dot{m}_{M,x_{1}/x_{1}+\Delta x}=\left( J_{I}A_{I} \right)_{x_{1}/x_{1}+\Delta x}$, where $A_{I}$ is the cross-sectional area of the interstitial volume biofilm and $J_{I}$ is the flux through this interstitial volume area. The mass-balances can be rewritten as

| $\frac{dm_{M}}{dt}=\left( J_{I}A_{I} \right)_{x_{1}}-\left( J_{I}A_{I} \right)_{x_{1}+\Delta x}-\dot{m}_{M,att}+\dot{m}_{M,mob}$ and | Eq. S3 |
| --- | --- |
| $\frac{dm_{A}}{dt}=\dot{m}_{M,att}-\dot{m}_{M,mob}.$ | Eq. S4 |

Both expressions can be re-written in terms of concentrations and concentration-based rates of attachment and mobilization via dividing by a volume. Rather than use total volume of the control volume, $V_{T}$, the interstitial volume within the control volume, $V_{I}$, will be used, harmonizing with the similar choice used for the flux expression. This yields the expressions

| $\frac{dc_{M,I}}{dt}=\frac{\left( J_{I}A_{I} \right)_{x_{1}}-\left( J_{I}A_{I} \right)_{x_{1}+\Delta x}}{V_{I}}-r_{M,att}+r_{M,mob}$ and | Eq. S5 |
| --- | --- |
| $\frac{dc_{A,I}}{dt}=r_{M,att}-r_{M,mob}$, | Eq. S6 |

which utilize the interstitial volume basis concentrations in the main text, and concentration-based rate quantities $r_{M,att/mob}$. To convert these expressions to a differential form, the volume term is decomposed to show $V_{I}=A_{I}\Delta x$ as $\Delta x\to0$. In addition, the cross-sectional area term is decomposed to show $A_{I}=\varphi A_{T}$ as $\Delta x\to0$ using the main text’s definition of porosity, $\varphi={V_{I}}/{V_{T}}$. Both relations are shown in a separate section. By substituting these equations, and taking the limit $\Delta x\to0$, the differential form of the mass-balances is found to be

| $\frac{\partial c_{M,I}}{\partial t}=\frac{1}{\varphi}\frac{\partial}{\partial x}\left( J_{I}\varphi\right)-r_{M,att}+r_{M,mob}$ and | Eq. S7 |
| --- | --- |
| $\frac{\partial c_{A,I}}{\partial t}=r_{M,att}-r_{M,mob}$*.* | Eq. S8 |

The interstitial-flux $J_{I}$ is then assumed to be dominated by Fickian diffusion, and thus can be replaced with Fick’s first law to show

| $\frac{\partial c_{M,I}}{\partial t}=\frac{1}{\varphi}\frac{\partial}{\partial x}\left( \varphi D_{B,I}\frac{\partial c_{M,I}}{\partial x} \right)-r_{M,att}+r_{M,mob}$ and | Eq. S9 |
| --- | --- |
| $\frac{\partial c_{A,I}}{\partial t}=r_{M,att}-r_{M,mob}$*.* | Eq. S10 |

Notably, because the interstitial flux was used, the interstitial concentration was used in the differential, along with an interstitial diffusivity in the biofilm $D_{B,I}$. By assuming that $D_{B,I}$ is constant throughout the biofilm, and expanding the differential,

| $\frac{\partial c_{M,I}}{\partial t}=D_{B,I}\left[ \frac{\partial^{2}c_{M,I}}{\partial x^{2}}+\frac{\varphi^{'}}{\varphi}\frac{\partial c_{M,I}}{\partial x} \right]-r_{M,att}+r_{M,mob}$ and | Eq. S10 |
| --- | --- |
| $\frac{\partial c_{A,I}}{\partial t}=r_{M,att}-r_{M,mob}$ | Eq. S11 |

are obtained. The mobilization and attachment rate terms can be defined using the same functional form presented by van Rossem *et al* (1) to be

| $r_{M,att}=k_{M,I}c_{M,I}c_{U,I}$ and | Eq. S12 |
| --- | --- |
| $r_{M,mob}=k_{A,I}c_{A,I}$*,* | Eq. S13 |

where $c_{U,I}$ is the concentration of unoccupied attachment sites within the control volume on an interstitial volume-basis. The concentration of open attachment sites can be replaced by the difference between the total number of attachment sites and the number of occupied attachment sites, $c_{U,I}=c_{S,I}-c_{A,I}$, to give the mass-balances

| $\frac{\partial c_{M,I}}{\partial t}=D_{B,I}\left[ \frac{\partial^{2}c_{M,I}}{\partial x^{2}}+\frac{\varphi^{'}}{\varphi}\frac{\partial c_{M,I}}{\partial x} \right]-k_{M,I}c_{M,I}\left( c_{S,I}-c_{A,I} \right)+k_{A,I}c_{A,I}$ and | Eq. S14 |
| --- | --- |
| $\frac{\partial c_{A,I}}{\partial t}=k_{M,I}c_{M,I}\left( c_{S,I}-c_{A,I} \right)-k_{A,I}c_{A,I}$*.* | Eq. S15 |

These mass-balances can then be directly converted to the governing equations presented in the main text using the dimensionless group definitions defined in the main text. This expression in nearly equivalent to the Fick-Jacobs expression per Dorfman and Yariv (2), with the exception of incorporating the expression $A_{I}=\varphi A_{T}$, allowing for the conversion to porosity, which is more prevalent in the biofilms literature.

**Showing** $\boldsymbol{V}_{\boldsymbol{I}}\boldsymbol{=}\boldsymbol{A}_{\boldsymbol{I}}\boldsymbol{\Delta x}$ **as** $\boldsymbol{\Delta x\to0}$ **and** $\boldsymbol{A}_{\boldsymbol{I}}\boldsymbol{=}\boldsymbol{\varphi}\boldsymbol{A}_{\boldsymbol{T}}$ **as** $\boldsymbol{\Delta x\to0}$**.**

The relation between the interstitial volume and interstitial cross-sectional area within the biofilm control-volume near the limit of $\Delta x\to0$ begins using the definition of a volume,

$V_{I}(x)=\int_{x_{1}}^{x_{1}+\Delta x} A_{I}(x)dx$,

Eq. S16

where the cross-sectional area is assumed to vary with depth throughout the biofilm, $A_{I}=f(x)$. To evaluate this integral, a functional form of the $A_{I}$ need be substituted. To do this, a Taylor series expansion on the function $A_{I}$ about $x_{1}$ is performed, yielding

$A_{I}\left( x \right)=A_{I}\left( x_{1} \right)+{A_{I}}^{'}\left( x_{1} \right)*\left( x-x_{1} \right)+\frac{{A_{I}}^{''}\left( x_{1} \right)}{2}*\left( x-x_{1} \right)^{2}+\ldots$.

Eq. S17

Given $\Delta x=x-x_{1}$, as $\Delta x\to0$, terms above order 2 become vanishingly small, leaving the expression

$A_{I}\left( x \right)=A_{I}\left( x_{1} \right)+{A_{I}}^{'}\left( x_{1} \right)*\left( x-x_{1} \right)$.

Eq. S18

Substituting this equation into the integral for $V_{I}$ eventually yields

$$V_{I}(x_{1})=A_{I}\left( x_{1} \right)*\Delta x$$

Eq. S19

as $\Delta x\to0$. This expression says that at point about which the control volume is taken, $x_{1}$, as the control volume shrinks, the interstitial volume of the shrinking control volume approaches the product of the cross-sectional area at the point of interest $x_{1}$ and the remaining control volume depth $\Delta x$. Considering this is the limit used to convert to the differential form, the interstitial volume can similarly be substituted using relation Eq. S19.

Porosity and cross-sectional area can be related using the definition of porosity for the considered control volume

$\varphi=\frac{V_{I}}{V_{T}}$.

Eq. S20

Using the definition $V_{T}=$ $\Delta xA_{T}$ and $V_{I}=A_{I}\Delta x$, it then follows that

$\varphi=\frac{A_{I}}{A_{T}}$

Eq. S21

only in the limit $\Delta x\to0$, which is necessary to satisfy Eq. S19.
